# piikun: An Information Theoretic Toolkit for Analysis and Visualization of Species Delimitation Metric Space

**DOI:** 10.1101/2023.08.02.551747

**Authors:** Jeet Sukumaran, Marina Meila

## Abstract

**Background:** Existing software for comparison of species delimitation models do not provide a (true) metric or distance functions between species delimitation models, nor a way to compare these models in terms of relative clustering differences along a lattice of partitions.

**Results:** `piikun` is a Python package for analyzing and visualizing species delimitation models in an information theoretic framework that, in addition to classic measures of information such as the entropy and mutual information [1], provides for the calculation of the variation of information criterion [2], a true metric or distance function for species delimitation models that is aligned with the lattice of partitions.

**Conclusions:** `piikun` is available under the MIT license from its public repository ( https://github.com/jeetsukumaran/piikun), and can be installed locally using the Python package manager ‘pip’.

## 1 Background

The field of species delimitation – computational approaches to determining the fundamental units of nature and atomic units of analysis in fields such as evolutionary biology and phylogenetics – has rapidly advanced in a number of ways recently, including machine learning approaches [3], incorporation of natural history [4], hierarchically modeling the speciation process over a primary population-level structuring [5] as well as other biological criteria [6]. Despite these advances, we still lack a true metric space to compare and relate various species delimitation models. Some useful indexes described previously, such as the taxonomic index of congruence (*C*_*tax*_) of [7] or the match ratio of [8] (implemented in [9]), are not true metrics (e.g., having the property of triangle inequality [10]), while others are defined for cases of nested models [7]. These limitations make intuitive interpretation of the distances challenging when more than two species delimitation models are involved, as well as less ideal in supporting inference or optimization algorithms in machine learning [2].

## 2 Implementation

This paper describes `piikun`^1^, a pure Python package [12] for the analysis and visualization of species delimitation models in an information theoretic framework that provides a true distance or metric space for these models. The package is publically available for download or local installation using ‘pip’ from its GitHub website https://github.com/jeetsukumaran/piikun, and depends on the following libraries: `NumPy` [13], `SciPy` [14], `PANDAS` [15, 16], `plotly` [17], `Matplotlib` [18], `Seaborn` [19].

The models analyzed using `piikun` may be generated by any inference package, such as BPP [20], DELINEATE [5] etc., or taxonomies or classifications based on conceptual descriptions in literature, geography, speculation, etc. Regardless of source or basis, each of these ways to organizae or cluster a set of lineages into a set of higher-level units is a (set theoretic) *partition* of those lineages [5, 21], and can be described in numerous ways that `piikun` can read (e.g., a generic JSON dictionary, or the a species delimitation model data exchange format “`SPART-XML`” [21]). `piikun` further supports specialized input formats, such as the comprehensive results file from DELINEATE [5] or BPP [20] analysis, which allow for incorporation of additional information, such as support values, as shown below.

`piikun` provides a range of univariate information theory statistics for each individual model in the input set (e.g., the entropy [1]), as well as bivariate statistics (e.g., the mutual information, joint entropy, [1]) for each distinct pair of these models, as well as *true* metrics (distances) between every pair of species delimitation models based on these information theoretic measures: the variation of information [2] and the normalized joint variation of information distance [22].

### 2.1 The Variation of Information Partition Distance

Every species delimitation model is a *partition* of a set of lineages into a set of mutually exclusive and jointly comprehensive subsets [21]. As such, the *variation of information* criterion of [2], which provides a true distance function for partitions, also establishes a metric space for species delimitation models. Given two partitions, *ψ*^*u*^, *ψ*^*v*^, *V I*(*ψ*^*u*^, *ψ*^*v*^), this is defined as [2]:

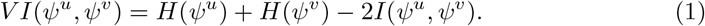

*H*(*ψ*) is the entropy of partition *ψ* which divides *n* elements into *K* subsets, with the *k*^*th*^ subset having *n*_*k*_ elements, and is given by [2]:

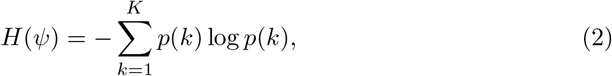

where *p*(*k*) is the probability of subset or cluster *k*, which is given in this approach by the cardinality of the subset as a proportion of the entire set: 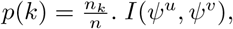. *I*(*ψ*^*u*^, *ψ*^*v*^), on the other hand, is the mutual information of partitions *ψ*^*u*^ and *ψ*^*v*^, and is given by:

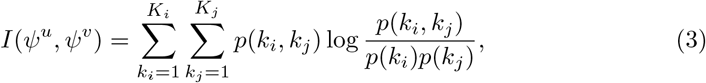

where *p*(*k*_*i*_, *k*_*j*_) is the joint probability of subset 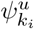 from partition *ψ*^*u*^ and subset 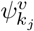 from partition *ψ*^*v*^. This is given by the proportion of the size of the intersection of subsets to the entire dataset: 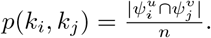.

### 2.2 Usefulness as a Species Delimitation Partition Metric

The information theoretic-based distances provided by `piikun` are very flexible. For example, different organizations of different populations into sets of species may be analyzed together from multiple different inferences or publications, even if the numbers of individuals, populations, or genes sampled across these sources vary. Furthermore, there are no constraints on the relationships between the partitions considered, such as being nested or otherwise. Detailed discussion of the statistical characteristics and properties of this statistic are given in [2]. Here we present a conceptual description of the properties of this metric that make it useful for analysis of differences between species delimitation models.

A true metric or distance function, such as the variation of information, has the properties of non-negativity, symmetry, and the triangle equality [10]. These properties are useful in aligning with human intuition when interpreting values, as well as preferable in statistical or computational terms due to benefits in algorithm and data structure design or scaling up comparisons [2].

In addition to being a true metric, the variation of information is aligned with the lattice of the set of partitions. Formally, a partition lattice is an ordered set of partitions, where the order is defined by the refinement of the partitions. We can represent the set of all partitions of a particular set of elements as a *partition lattice* using the refinement order (a partition *U* is defined as a refinement of a partition *V* if every block in *U* is a subset of a block in *V* : we say that *U* is finer than *V*, or, equivalently, *U* is coarser than *V* ). Conceptually, a partition lattice provides geometrical representation of all possible divisions of a set of elements, from the most granular, where each element forms its own subset, to the most general, where all elements belong to a single subset. This allows for the understanding of relationships between different partitions, offering insights into how small changes in groupings can lead to new partitions, and how these partitions are (or are not) nested within each other. A metric on partitions that is aligned along the lattice of partition sets has numerous advantages for interpretation of the disagreements between different partitions as well as for computation as neighbors in the metric space correspond to refinements in the partition lattice [23]. Such a metric takes into account the hierarchical nature of partitions, recognizing that partitions related through a series of refinements are closer to each other than partitions that differ more fundamentally. That is, under this metric, the nearest neighbor of a species delimitation partition can be obtained by “lumping” the two smallest species blocks of that partition, and vice versa via “splitting”. This alignment with the partition lattice is not only useful for intuition, but may also have potential to provide benefits in designing inference or stochastic sampling algorithms (e.g., MCMC moves in partition space).

The upper bound of this metric is given by 2 log *K*, where *K* is the maximum number of subsets in a partition, or by *log*(*n*), with *n* the number of elements (populations or individuals, depending on the approach), if any partition is possible [2]. In some applications, it might be useful to have this normalized to the range [0, 1]. Unfortunately, while it is straight-forward enough to normalize by these known upper bounds, this will preclude comparison of distances across different datasets (in particular, datasets of different sizes). For this latter purpose, `piikun` provides for the calculation of the normalized variation of information following [22], which retains the metric properties of the variation of information.

## 3 Results: Example Exploratory Discovery Analysis

Here we focus on one of the visualizations offered by `piikun` which gives us insight into how the “disagreement” between two partitions, as measured by their distance, might vary with respect to some value, trait, or attribute of each of the respective partitions. The visualization in Figure 1 shows how different species delimitation partition distances might vary in relation to the support (probability) of each partition in the comparison. `piikun` was run on the 1000 most probable species delimitation models from a DELINEATE inference on a dataset of *Lionepha* beetles [24, available on the DELINEATE website]. The partitions with high probabilities (i.e., the upper-right area of Figure 1) are mutually close. In contrast, the less probable partitions are much more dissimilar, pairwise (i.e., the lower-left area of Figure 1), while remaining relatively similar to the group of probable partitions (i.e., the right edge of Figure 1). This strongly suggests the existence of a compact “core” formed of the most probable partitions, surrounded by a “halo” of diverse, less probable partitions. Further analysis (e.g., explicit enumeration and comparison of the subsets of the partitions) could then be used confirm that this is indeed because the species delimitation partitions with higher probabilities are disagreeing on the finer-scale splitting of species, while the less probable models, conflict with each other in more fundamental ways. Other datasets might indicate other relationships, e.g. where more supported models show more fundamental differences from one another, serving as the motivation or bases for more directed statistical analysis at the identified regions of partition space of interest.

**Fig. 1.**
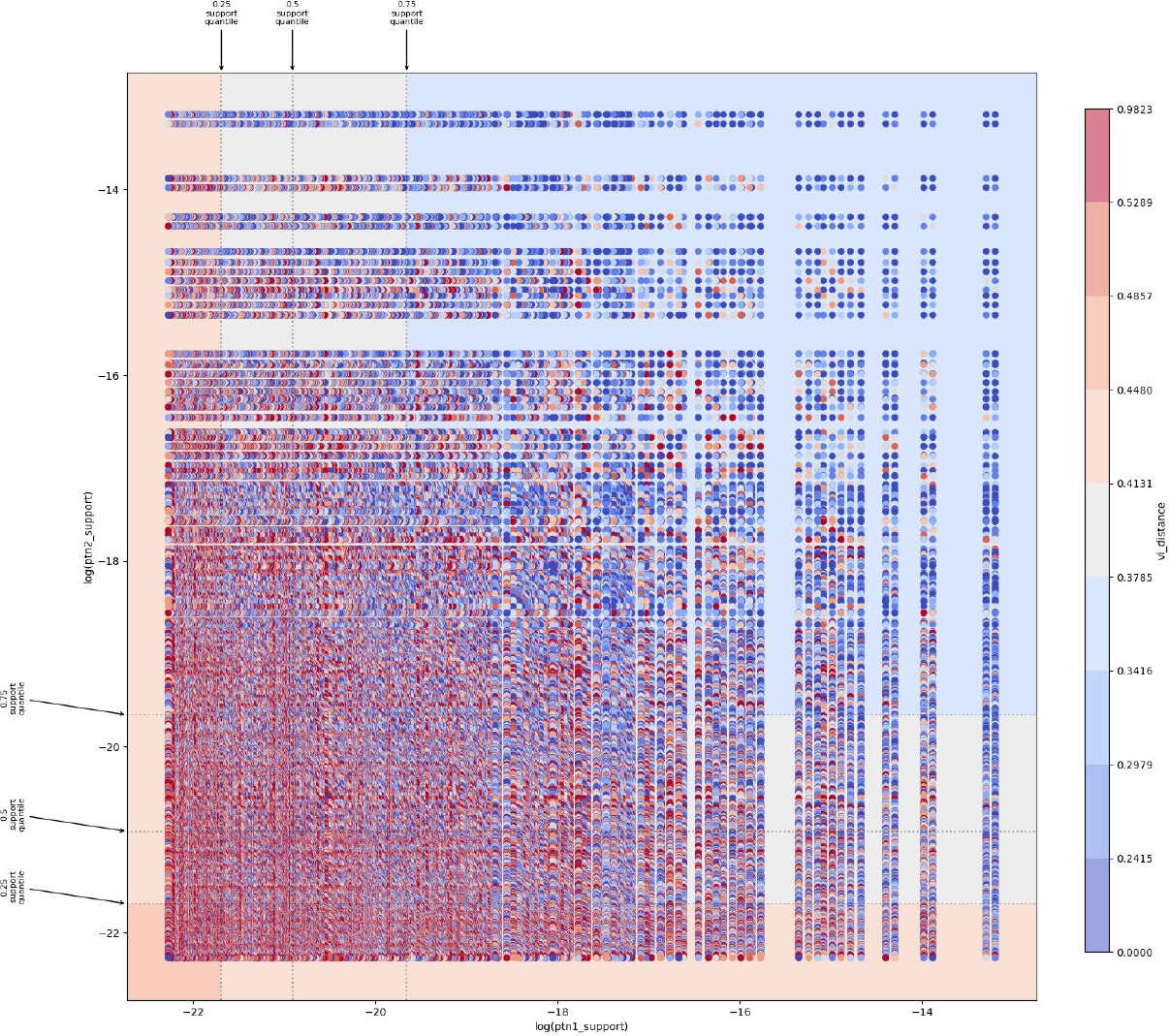
Understanding how the pairwise similarity of competing species delimitation models varies with their probability (support). The log(`ptn_1`) and log(`ptn_1`) axes represent species delimitation models (partitions) ordered by their log-scaled probabilities. Each point represents a pair of models; its color encodes the variation of information distances between them. The color gradient goes from blue to red, with blue indicating smaller distances while red indicating greater distances. The plot background is regionalized into 0.25, 0.5, and 0.75 quantiles based on support, with mean distance for partitions compared within each region indicated with the same color scheme. Note how partitions with high probabilities (upper-right area) are mutually close (VI-distance almost 0), while partitions with lower probabilities (lower-left area) are much more different (VI-distance over 0.91).

## 4 Conclusions

`piikun` is a Python package of command-line tools for generating insight about *differences* between species delimitation models, beyond ranking them in terms of quality of inference or data.

`piikun` provides a general implementation of the variation of information criterion [2], a metric function returning distances between partitions of a set, that, in addition to being a metric has a range of particular characteristics that are especially useful when used to establish a metric space for species delimitation models, such as alignment along the lattice of partitions, which supports intuitive interpretation of the differences with reference to nesting of the models. As with all information theoretic approaches, the metric extracts signal from patterns in the data without reference to any mechanistic or phenomenological processes or assumptions, which allows species delimitation models to be compared across inferrential models and datasets. This combination of being a true metric, alignment along the lattice of partitions, and being able to compare not just species delimitation models, but essentially any partitioning of elements as long as the concepts can be represented as an arbitrary nesting of arbitrary labels expressible as a JSON list (with more specialized formats for associationg metadata), allows for the application of `piikun` to provide insight into differences in domains well beyond systematics or even biology. With documentation that works the user through a complete workflow, a modular design for UNIX-pipeline style composibility, and the ability to be run directly without explicit userspace installation using ‘pipx’ or easy local installation using ‘pip’, this utility will facilitate research in species development modeling as well as make a wider range of post-inferential analyses and visualization more accessible to empirical researchers in evolutionary biology and related fields.

## 5 Availability and requirements

**Project name** `piikun`

**Project home page** https://github.com/jeetsukumaran/piikun

**Operating system(s)** Platform independent

**Programming language** Python

**Other requirements** Python 3.10 or higher

**License** New BSD License

## 6 Declarations

### 6.1 Funding

This work was possible due to support from the National Science Foundation grant to author JS: NSF-DEB 1937725 “COLLABORATIVE RESEARCH: Phylogenomics, spatial phylogenetics and conservation prioritization in trapdoor spiders (and kin) of the California Floristic Province”.

### 6.2 Conflict of interest/Competing interests

None.

### 6.3 Ethics approval and consent to participate

Not applicable.

### 6.4 Consent for publication

Not applicable.

### 6.5 Data availability

- https://github.com/jeetsukumaran/piikun
- https://drive.google.com/file/d/1GQz55ngL1-slg4sCCMuWcnTYA3XADDmc/view?usp=drive_link

### 6.6 Materials availability

Not applicable.

### 6.7 Code availability

https://github.com/jeetsukumaran/piikun

### 6.8 Author contribution

JS developed and is responsible for maintaining the software, applying theory and fundamental expressions of information in partition space previously developed by MM [2] to species delimitation models and analyses in evolutionary biological space. MM, in addition to development of the original variation of information criteria theory that is the theoretical grounding of this software, provided conceptual, qualitative and quantitative test-based oversight of the software itself.

Otherwise, both authors contributed equally in all other respects to this work, including the writing, experimental design, and approaches to analysis and visualization.

## Footnotes

1 “sparrowhawk” in the Kumeyaay language [11]. San Diego State University is built on Kumeyaay land.

## Notes

https://github.com/jeetsukumaran/piikun

### Competing Interest Statement

The authors have declared no competing interest.

### Summary of Updates

Revised/reformatted/restructured for new submission to journal.

